# CARMIL3 is important for cell migration and morphogenesis during early development in zebrafish

**DOI:** 10.1101/2020.09.27.315655

**Authors:** Benjamin C. Stark, Yuanyuan Gao, Lakyn Belk, Matthew A. Culver, Bo Hu, Diane S. Sepich, Marlene Mekel, Lilianna Solnica-Krezel, Fang Lin, John A. Cooper

## Abstract

Cell migration is important during early animal embryogenesis. Cell migration and cell shape are controlled by actin assembly and dynamics, which depend on capping proteins, including the barbed-end heterodimeric actin capping protein (CP). CP activity can be regulated by capping-protein-interacting (CPI) motif proteins, including CARMIL (capping protein Arp2/3 myosin-I linker) family proteins. Previous studies of CARMIL3, one of the three highly conserved CARMIL genes in vertebrates, have largely been limited to cells in culture. Towards understanding CARMIL function during embryogenesis *in vivo*, we analyzed zebrafish lines carrying mutations of *carmil3*. Maternal-zygotic mutants show impaired endodermal migration during gastrulation, along with defects in dorsal forerunner cell (DFC) cluster formation, affecting the morphogenesis of Kupffer’s vesicle (KV). Mutant KVs are smaller and display decreased numbers of cilia, leading to defects in left/right (L/R) patterning with variable penetrance and expressivity. The penetrance and expressivity of the KV phenotype in *carmil3* mutants correlated well with the L/R heart positioning defect at the end of embryogenesis. This first *in vivo* animal study of CARMIL3 reveals its new role for CARMIL3 during morphogenesis of the vertebrate embryo. This role involves migration of endodermal cells and DFCs, along with subsequent morphogenesis of the KV and L/R asymmetry.

## Introduction

### CARMIL Regulation of Actin Assembly via Capping Protein

CARMILs are one family of capping-protein-interacting (CPI-motif) proteins, reviewed in (Edwards et al., 2014). Vertebrates, including zebrafish, have three conserved CARMIL-encoding genes, called *CARMIL1*, *CARMIL2* and *CARMIL3* in humans (Stark et al., 2017; Stark and Cooper, 2015). The zebrafish genes, *carmil1, carmil2* and *carmil3,* have distinct spatial and temporal expression patterns during development (Stark and Cooper, 2015). In human cultured cells, available evidence indicates that the gene products have distinct subcellular locations and functions, even within one cell type (Lanier et al., 2015; Liang et al., 2009; Stark et al., 2017). The functions of *CARMIL1* and *CARMIL2* include cell migration, macropinocytosis, lamellipodial activity and cell polarity.

In contrast to this information for CARMIL1 and CARMIL2, relatively less is known about the *CARMIL3* gene. In mouse, the CARMIL3 protein localizes to developing synapses and spines in neurons, where it recruits actin capping protein. Depletion of CARMIL3 protein in neurons leads to defects in spine and synapse assembly and function (Spence et al., 2019). In breast and prostate cancer patients, elevated expression of *CARMIL3* correlates with poor outcomes, and mouse tumor models reveal a role for *CARMIL3* in epithelial-mesenchymal transition, cadherin-based cell adhesions, and cell migration and invasion (Wang et al., 2020).

### Cell Migration and Morphogenesis in Early Vertebrate Development

Early vertebrate embryogenesis sees massive cell rearrangements that establish and shape the three germ layers, mesoderm, endoderm and ectoderm during gastrulation. The most deeply positioned endoderm gives rise to the gut and other alimentary organs. At the onset of zebrafish gastrulation, mesendodermal progenitors are located at the margin of a cup-shaped blastoderm that covers the animal hemisphere of a large yolk cell (Warga and Kimmel, 1990). The mesodermal and endodermal lineages soon separate, and internalized endodermal cells initially disperse on the yolk surface as individuals via a random walk to almost completely cover the yolk cell (Pézeron et al., 2008). Concurrently, mesoderm and ectoderm spread around the yolk cell in the process of epiboly (Warga and Kimmel, 1990). Later during gastrulation, endoderm cells migrate towards the dorsal midline along trajectories that are biased either animally/anteriorly (for cells in animal hemisphere) or vegetally/posteriorly (for cells in vegetal hemisphere), thus simultaneously elongating the nascent endoderm along the AP axis (Schmid et al., 2013). Endodermal cell migration during gastrulation depends on Rac1-regulated actin dynamics (Woo et al., 2012).

On the dorsal side there is a small cluster of dorsal forerunner cells (DFCs) that travel vegetalward in advance of the spreading germ layers, which later during segmentation will form an epithelial ciliated vesicle known as Kupffer’s Vesicle (KV), the left-right (L/R) organizer of zebrafish (Amack and Yost, 2004). At the end of epiboly, DFCs form multiple rosette-like epithelial structures whose focal points are enriched for apical proteins. During segmentation, these rosettes arrange into a single rosette lined by a lumen with cilia at the apical membrane of the cells, thereby forming the KV (Oteíza et al., 2008). L/R patterning in zebrafish depends on the motile cilia in the KV that generate an asymmetric fluid flow (Gokey et al., 2015; Gokey et al., 2016; Sampaio et al., 2014). Mutations that affect the shape and size of the KV or that affect the number or length of cilia in the KV can impair robust L/R patterning (Amack, 2014).

To investigate the function of CARMIL3 in vertebrate development, we examined phenotypes resulting from the disruption of the gene encoding CARMIL3 in zebrafish. We found defects in endodermal cell migration, DFC migration and clustering, KV morphogenesis, the number of cilia in the KV, and L/R asymmetry.

## Materials & Methods

### Zebrafish lines and husbandry

Animal protocols were approved by the Institutional Animal Care and Use Committees at University of Iowa and Washington University. At University of Iowa, zebrafish were maintained as described previously (Xu et al., 2011) and embryos were obtained by natural spawning and staged according to morphological criteria or hours post fertilization (hpf) at 28°C or 32°C unless otherwise specified, as described previously (Kimmel et al., 1995). At Washington University, zebrafish and embryos are maintained at 28.5°C using the standard operating procedures and guidelines established by the Washington University Zebrafish Facility, described in detail at http://zebrafishfacility.wustl.edu/documents.html. The following zebrafish lines were used in this study: AB*/Tuebingen, *Tg(sox17*:*EGFP)* (Mizoguchi et al., 2008), *carmil3*^*sa19830*^ (*lrrc16b*^*sa19830*^), and *carmil3*^*stl413*^.

Zebrafish line *carmil3*^*sa19830*^ (*lrrc16b*^*sa19830*^) was obtained from the Zebrafish International Resource Center (Eugene, OR) (described at https://zfin.org/ZDB-ALT-131217-14950) (Kettleborough et al., 2013). The mutation was created by *N*-ethyl-*N*-nitrosourea (ENU) treatment of adult males, and the mutated gene has a G to T conversion at an essential splice site of intron 27-28, which introduces multiple stop codons beginning at amino acid residue 832 (Kettleborough et al., 2013). Failure to splice at this site is predicted to change amino-acid residue 831 from E to D, with the next codon being ochre TAA, and thus truncating the protein to 832 residues from its normal length of 1384 residues. Zebrafish line *carmil3*^*stl413*^ was produced via TALEN-mediated mutagenesis (Boch et al., 2009; Moscou and Bogdanove, 2009). TALEN sequences used were 5’-TGACAAGACATCAATCAAGT and 5’-TTTGCCACTCTTGTTCTCTG (corresponding to bases 72063-72082 and 72101-72120, respectively, of the sequence for the genomic locus FQ377660). TALEN cleavage led to several different independent mutations in founder fish. The largest deletion was of an 11-bp segment of exon 2, 5’-ACGTATCAAAG. This deletion eliminates the Alu1 restriction site found in exon 2, and the deletion leads to a premature stop codon at amino-acid residue 101. Fish carrying this mutation were selected for outcrossing and further study.

*carmil3*^*sa19830*^ and *carmil3*^*stl413*^ were genotyped by restriction enzyme digestion of PCR amplicons containing the mutations. For *carmil3*^*sa19830*^ the forward primer was 5’-AGCAGAGTGTCTTTCTCCAC, and the reverse primer was 5’-GATCGAGGTTGGAGGTGAAC. MseI digest distinguished WT (191 bp band) from *sa19830* mutant (bands at 123 bp and 38 bp). For *carmil3*^*stl413*^ the forward primer was 5’-AGAATAGTGTAATCCACTCATTTTTCAACCG, and the reverse primer was 5’-AGGCAGGTGTGAATACCTTTAAAGTCTTCA (corresponding to bases 71935-71965 and 72249-72278, respectively, of the sequence for the genomic locus FQ377660). AluI digestion produced two bands at 157 bp and 187 bp from WT genomic DNA, and a single band at 333 bp from the mutant. Mutant founders were outcrossed into WT AB fish to produce heterozygous fish, which were fully viable, and they were mated to produce maternal and maternal-zygotic homozygous mutants. Alternatively, genotyping of *carmil3*^*sa19830*^ mutants was performed by Transnetyx (Cordova, TN) using real-time PCR with allele-specific probes.

### Protein expression and purification

Capping protein (CP, mouse alpha1beta2) was expressed and purified as described (Johnson et al., 2018). Glutathione-S-transferase (GST)-tagged CP binding region (CBR) fragments of human CARMIL1a (E964-S1078, plasmid pBJ 2411), human CARMIL3 (S955-S1063, plasmid pBJ 2449), zebrafish Carmil3 (S943-N1040, plasmid pBJ 2451) and zebrafish Carmil3 CPI-mutant (S943-N1040, with point mutations H944A and R966A, plasmid pBJ 2452) were expressed from pGEX-KG vectors in *E. coli* BL21 Star (DE3). The fusion proteins were affinity-purified on Glutathione Sepharose® 4 Fast Flow (GE Healthcare), and then bound to POROS GoPure XS (Applied Biosystems) in 20 mM NaH_2_PO_4_, 100 μM EDTA, 1 mM NaN3, 5 mM DTT, 4M urea (pH 7.5). After elution with a gradient to 1 M NaCl in the same buffer, the purified GST-CBR fragments were concentrated and stored at −70°C.

### Actin polymerization assays

Pyrene-actin polymerization assays were performed as described (Carlsson et al., 2004). Pyrene-labeled and unlabeled gel-filtered rabbit muscle actin stocks were mixed to produce a total actin monomer concentration of 1.5 μM in the cuvette. Pyrene-actin filament seeds were prepared as described (Ramabhadran et al., 2012). CP at 5 nM and GST-CBR at varied concentrations were added at the start of the experiment (0 sec). Pyrene-actin fluorescence was measured using time-based scans on a steady-state fluorometer (QuantaMaster, PTI, Edison, NJ) with excitation at 368 nm and emission at 386 nm.

### Whole-mount RNA in situ hybridization (WISH)

Digoxigenin-labeled antisense RNA probes for *myl7* (cardiac myosin regulatory light chain) (Ye and Lin, 2013; Yelon et al., 1999), *sox17* (sex determining region Y-box 17) (Alexander et al., 1999; Hu et al., 2018), *southpaw* (*spaw*) (Long et al., 2003; Panizzi et al., 2007) were synthesized by *in vitro* transcription. Staged embryos were fixed in 4% fish fix solution (4% paraformaldehyde, 4% sucrose, 0.1 M phosphate buffer pH 7.2, 0.12 mM CaCl_2_) at 4°C overnight. Fixed embryos were manually dechorionated and dehydrated with a series of methanol washes. WISH was performed as described (Thisse and Thisse, 2008).

### Whole-mount Immunofluorescence

Staged embryos (10-14 somites) were manually dechorionated and fixed in Dent’s fixative (80% methanol: 20% DMSO) at room temperature for a minimum of 2 hrs. Antibody staining was performed in PBDT (1% BSA, 2% goat serum, 2% DMSO, 0.1% Triton X-100 in PBS) as described (Topczewski et al., 2001; Ye and Lin, 2013). The following antibodies were used: mouse anti-acetylated tubulin (clone 6-11b-1, Sigma-Aldrich, diluted 1:2000) and goat anti-mouse Alexafluor 488 (Invitrogen, diluted 1:2000).

### Microscopy and Image Analysis

WashU: For Figure 4 D-G, fluorescence images were collected on a spinning disk confocal microscope (Quorum, Canada) using an inverted Olympus IX-81 microscope, a Hamamatsu EMCCD camera (C9100-13) and Metamorph acquisition software. For Figures 2A, 5, and 6, WISH images were collected on a Nikon Macroscope with a Nikon AZ100 objective, a 1x lens N.A. 0.1, and a 4x lens N.A. 0.4.

Iowa: For still epifluorescence images, live or fixed embryos were mounted in 2% methylcellulose and photographed using a Leica DMI 6000 microscope with a 5×/NA 0.15 objective or a 10×/NA 0.3 objective. For ISH images in Figure 3, embryos were mounted in 80% glycerol/PBS and photographed using a Leica M165FC Stereomicroscope with a Leica DFC290 Color Digital Camera.

For time-lapse imaging of endodermal cells in Figure 2 C-G and Figure 4 A-C, *Tg(sox17:EGFP)* embryos were embedded in 0.8% low-melting agarose in a dorsal-mount imaging mold as previously described (Ye et al., 2015). Time-lapse imaging was taken in the dorsal region of endoderm at 25°C, at 5-minute interval with a 5×/NA 0.15 objective on an inverted Leica DMI 6000 microscope. Images were processed and cell tracking analyzed using ImageJ (Schneider et al., 2012). Data were exported to Excel where cell migration speed, paths, direction were determined as previously reported (Lin et al., 2005).

### Statistical Analysis

Data were compiled from two or more independent experiments and were presented as the mean ± SEM or ± SD as indicated in figure legends. Statistical analyses were performed in GraphPad Prism (GraphPad Software) using unpaired two-tailed Student's *t*-tests with unequal variance. The numbers of cells and embryos analyzed in each experiment and significance levels are indicated in graphs and/or the figure legends.

## Results

As part of our interest in how actin assembly and actin-based motility contributes to morphogenesis and cell movement during development, we examined the role of CARMIL3. The CARMIL family of proteins regulate the heterodimeric actin capping protein (CP) that controls polymerization of actin filaments at barbed ends, *in vitro* and in cells (Stark et al., 2017). To advance our understanding of the functions of the three CARMIL isoforms conserved across vertebrates, we first assayed biochemical activities of zebrafish Carmil3 in comparison to human CARMIL3 and CARMIL1. Second, we altered the activity of the zebrafish *carmil3* gene, previously known as *lrrc16b* and *si:ch211-204d18.1*, by creating and/or interrogating lines carrying homozygous loss-of-function mutations.

### CARMIL3 inhibits actin capping activity of CP

The biochemical activities of the vertebrate CARMIL1 and CARMIL2 isoforms with respect to CP and actin have been studied in mouse and human systems (Stark et al., 2017). The ~100-aa capping protein binding region (CBR) of CARMIL1 and CARMIL2 are known to bind directly to CP and partially inhibit its actin capping activity, via an allosteric mechanism. Comparative studies of CARMIL3 had not been previously done and so we performed them here.

We prepared recombinant proteins corresponding to the CBR region of CARMIL3 from human and zebrafish, and we tested their ability to inhibit the capping activity of mouse non-sarcomeric CP (alpha1beta2) (Edwards et al., 2013). In actin polymerization assays seeded with barbed ends of actin filaments, CP inhibited actin polymerization (Fig. 1A). A human CARMIL3 (C3) CBR fragment partially reversed the inhibitory effect of CP, with a potency slightly greater than that of human CARMIL1 (C1) (Fig. 1A). Zebrafish Carmil3 (C3) CBR also inhibited CP (Fig. 1B), with a potency approximately half that of human CARMIL3. Specific point mutations known to impair the ability of human CARMILs to interact with CP (Edwards et al., 2013; Lanier et al., 2015) were also created in the CBR fragment of zebrafish Carmil3 (C3-HRAA). The mutant CBR failed to inhibit CP activity (Fig. 1 B). Thus, the CBR region of CARMIL3, from both human and zebrafish, is able to inhibit the capping activity of CP, in a manner consistent with the direct-binding mechanism described for other CARMILs (Stark et al., 2017). This result supports the hypothesis that functional properties of CARMIL3 in vertebrates may involve direct interaction with CP.

**Figure 1.**
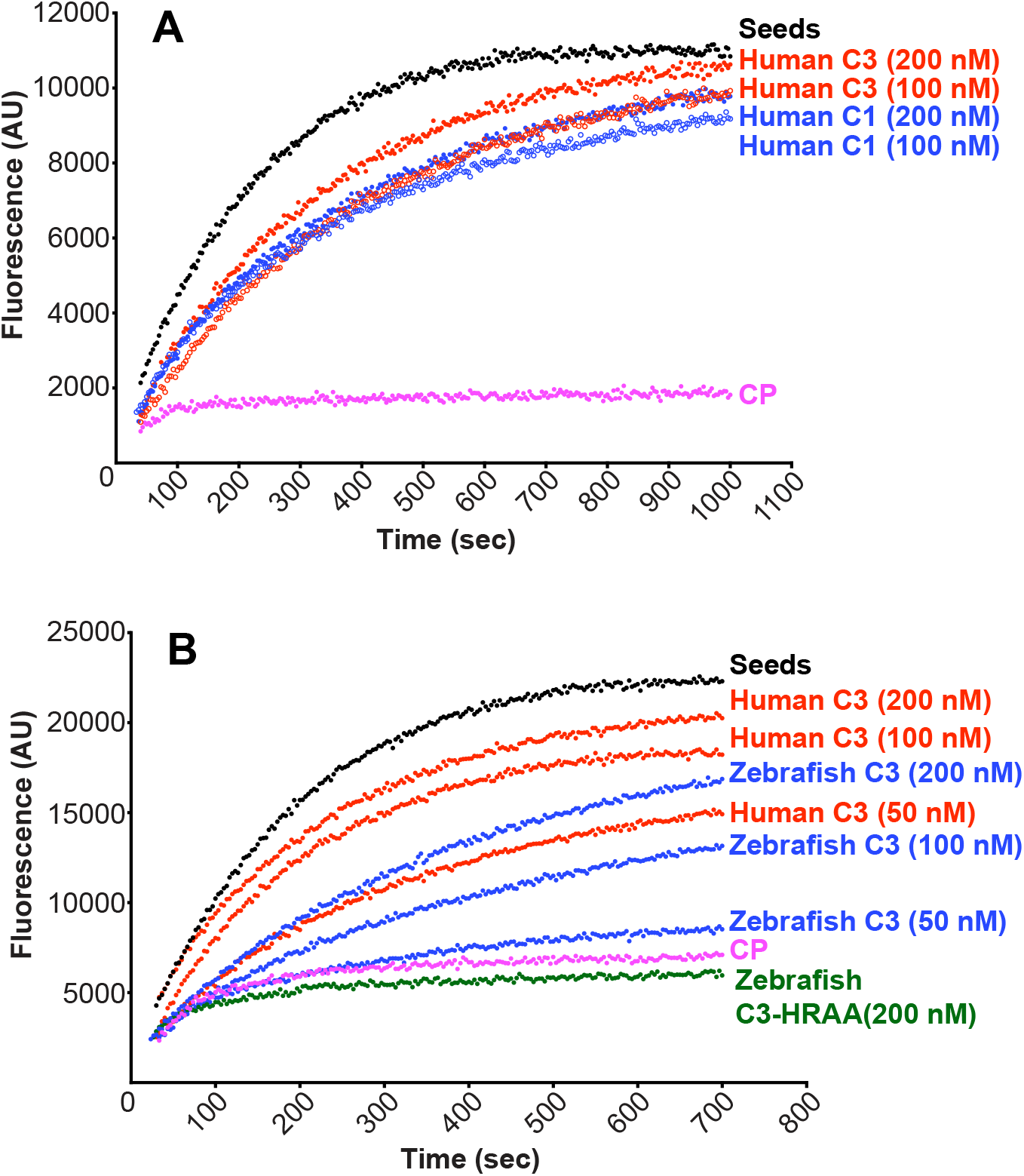
CARMIL3 inhibits capping protein (CP) activity in actin polymerization assays, with pyrene-actin fluorescence (arbitrary units) plotted vs time. A) Human CARMIL3 (C3) inhibited the ability of CP to cap barbed ends of actin filaments. Black points, labeled “Seeds,” are from a control with monomeric actin and filamentous actin seeds. Pink points, labeled “CP,” correspond to a sample to which CP was added. Red and blue curves contain CP plus the indicated concentrations of human CARMIL1 or CARMIL3 CBR fragment. B) Zebrafish CARMIL3 (C3, blue curves) inhibited capping by CP. In comparison to human CARMIL3 (C3, red curves), the inhibitory activity of zebrafish CARMIL3 was slightly less than that of human CARMIL3, based on similar concentrations as indicated. A mutant form of zebrafish CARMIL3 (C3-HRAA, green curve) containing two point mutations at conserved residues of the CP-binding CPI motif, failed to block CP activity, as expected.

### Genomic mutations of zebrafish carmil3: Creation and characterization

A *carmil3* mutant line from the Zebrafish Mutation Project (Kettleborough et al., 2013), *carmil3*^*sa19830*^, carries a single-nucleotide change at an essential splice site, predicted to truncate the protein to 832 residues from its WT length of 1384 residues. We created a second *carmil3* mutant line by TALEN-mediated mutagenesis, *carmil3*^*stl413*^; this mutation produces an 11-bp deletion within exon 2 that causes a frameshift followed by a nonsense mutation, predicted to produce a truncated protein of 101 residues. Heterozygous and homozygous zygotic mutants of both alleles were viable as embryos and adults and did not present any overt phenotypes. When fish homozygous for either of the two alleles were crossed, the resulting embryos, deficient in both maternal and zygotic *carmil3* function, MZ*carmil3*^*sa19830/sa19830*^ (thereafter MZ*carmil3*^*sa19830*^) or MZ*carmil3*^*stl413/413*^ (thereafter MZ*carmil3*^*stl413*^), also completed epiboly and gastrulation and developed into morphologically normal embryos.

### Migration of endodermal cells

To assess potential subtle or transient developmental defects, we analyzed the expression of germ layer markers in MZ*carmil3* mutants at mid-gastrulation by whole mount *in situ* hybridization (WISH). We examined the positions of endodermal cells and DFCs, marked by expression of *sox17*, over time during gastrulation. In embryos fixed and stained by whole-mount in situ hybridization (WISH) at midgastrulation (70-80% epiboly), we noted that the leading edge of the vegetally migrating endodermal cells in MZ*carmil3*^*sa19830*^ mutant embryos lagged behind that of WT embryos (Fig. 2A). To quantify the effect, we measured the distance from the front edge of the endodermal cells to the edge of the aggregation of DFCs, as diagrammed on the right side of Figure 2B. The distance was larger for the MZ*carmil3*^*sa19830*^ mutant, compared to WT embryos (Fig. 2B), indicating that *carmil3* is important for endodermal migration. The mean distance values were 67 μm (95% confidence interval of 60-74, n=61) for WT embryos and 142 μm (95% confidence interval of 135-150, n=55) for mutant embryos. The difference between the two sets of data had a two-tailed P value of <0.0001 in an unpaired t test. These data were combined from clutches obtained on separate days.

**Figure 2.**
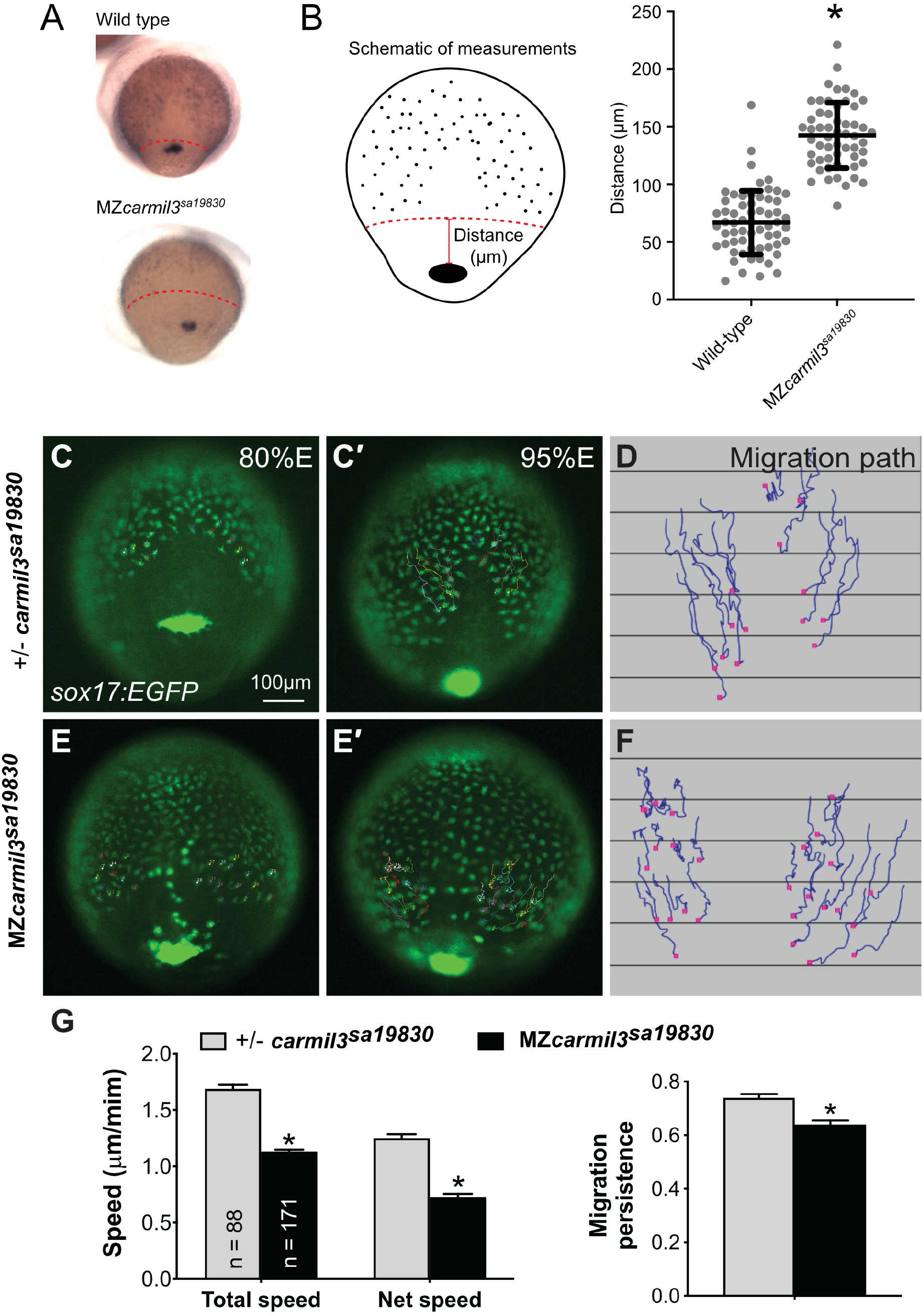
Pattern of endodermal cell migration in *carmil3* mutant embryos compared to WT embryos. Panels A and B are results from *sox17* staining of embryos at 70-80% epiboly. A. Representative images illustrating the patterns observed in WT compared to MZ*carmil3*^*sa19830*^ mutant embryos. Red dotted line indicates the margin of the endoderm, used to measure migration distance. B. Endodermal cell migration distribution, measured as illustrated in the schematic and the images of panel A. In the plot, each data point corresponds to one embryo. Values for the distributions were as follows (mean ± s.d.): WT 74 ± 25 (N=50), MZ*carmil3*^*sa19830*^ mutant 144 ± 28 (N=44). Asterisk (*) indicates p value of <0.0001 in Student’s *t-*test. Panels C through G are results from epifluorescence time-lapse experiments performed on MZ*carmil3*^*sa19830*^ or MZ*carmil3*^*sa19830*^ mutant embryos, each carrying *Tg(sox17:EGFP*)/+. (C and E) Snapshots at 80% epiboly stage from the time-lapse movie, with the tracked cells labeled. (C’ and E') Snapshots at 95% epiboly stage, with the migration tracks of endodermal cells from the 80% to 95% epiboly stage superimposed. Scale bar: 100μm. (D, F) Migration tracks delineate routes of endodermal cells. Solid magenta squares denote the endpoint of migration. (G) Total speeds, net speeds, and migration persistence. The number of cells analyzed is indicated in the first graph. Asterisk (*) indicates p value of <0.0001 in Student’s *t*-test.

To test this observation directly in living embryos, we then examined the migration of the endodermal cells by conducting fluorescence microscopy time-lapse analyses using a transgenic line expressing eGFP fluorescent protein from the *sox17* gene promoter Tg(*sox17-GFP*) (Fig. 2) (Mizoguchi et al., 2008). Cell tracking revealed that in control embryos (heterozygous *carmil3^sa19830/+^*), endodermal cells migrate toward the vegetal pole and converge dorsally along fairly straight paths (Schmid et al., 2013) (Fig. 2C-D). In contrast, in *carmil3*-deficient mutants (MZ*carmil3^sa19830^*) the endodermal cells took less direct paths (Fig. 2E-F). Further cell tracking analyses showed that both cell movement, i.e. (total speed: movements in all directions) and migration efficiency (net speed: along straight line between the start and endpoint) were impaired in mutants compared to WT controls, as was the persistence of migration (ratio of net: total speed) (Fig. 2G). Thus, Carmil3 is required for efficient migration of endodermal cells.

### Aggregation of dorsal forerunner cells (DFC)

Static images from these time-lapse experiments with a *Tg[sox17:EGFP]* background also revealed the migration and cluster formation by dorsal forerunner cells (DFCs), which strongly expressed eGFP. DFCs in WT embryos usually formed a single large cluster (Fig. 2C). In mutant embryos, the DFC cluster was fragmented; we observed this in MZ*carmil3*^*sa19830*^ embryos (Fig. 2E) and in MZ*carmil3*^*stl413*^ mutant embryos (data not shown). In addition, we noted bright GFP-labeled cells that resided in an apparent gap between the DFC cluster and the leading edge of the migrating endodermal cells (Fig. 2E) in the mutant, but not WT embryos, which we suggest correspond to DFCs exhibiting abnormal migration.

We further examined DFC cluster morphology and DFC distribution during gastrulation using WISH probes for cells expressing *sox17*; these results also revealed the location of DFCs over time in larger populations of WT and mutant embryos. We examined embryos at 70-80% epiboly (midgastrulation; Fig. 3), and we observed variation in the size and coherence of the DFC cluster in the mutants compared to WT. To score the phenotype, we graded the morphology of the aggregated set of DFCs as “normal size, one cluster;” “normal size, split-mild,” “normal size, split-severe;” “small size, one cluster,” and “small size, split-mild.” Representative images are shown in Figure 3A-D, and graphs with quantification of these results, for control, MZ*carmil3*^*sa19830*^, and *carmil3*^*stl413*^ mutant embryos, are presented in Fig. 3E. These results corroborate the observations from the time-lapse analyses, showing that formation of a cohesive DFC cluster is impaired in *carmil3*-deficient gastrulae, which frequently displayed a fragmented and/or smaller DFC aggregate.

**Figure 3.**
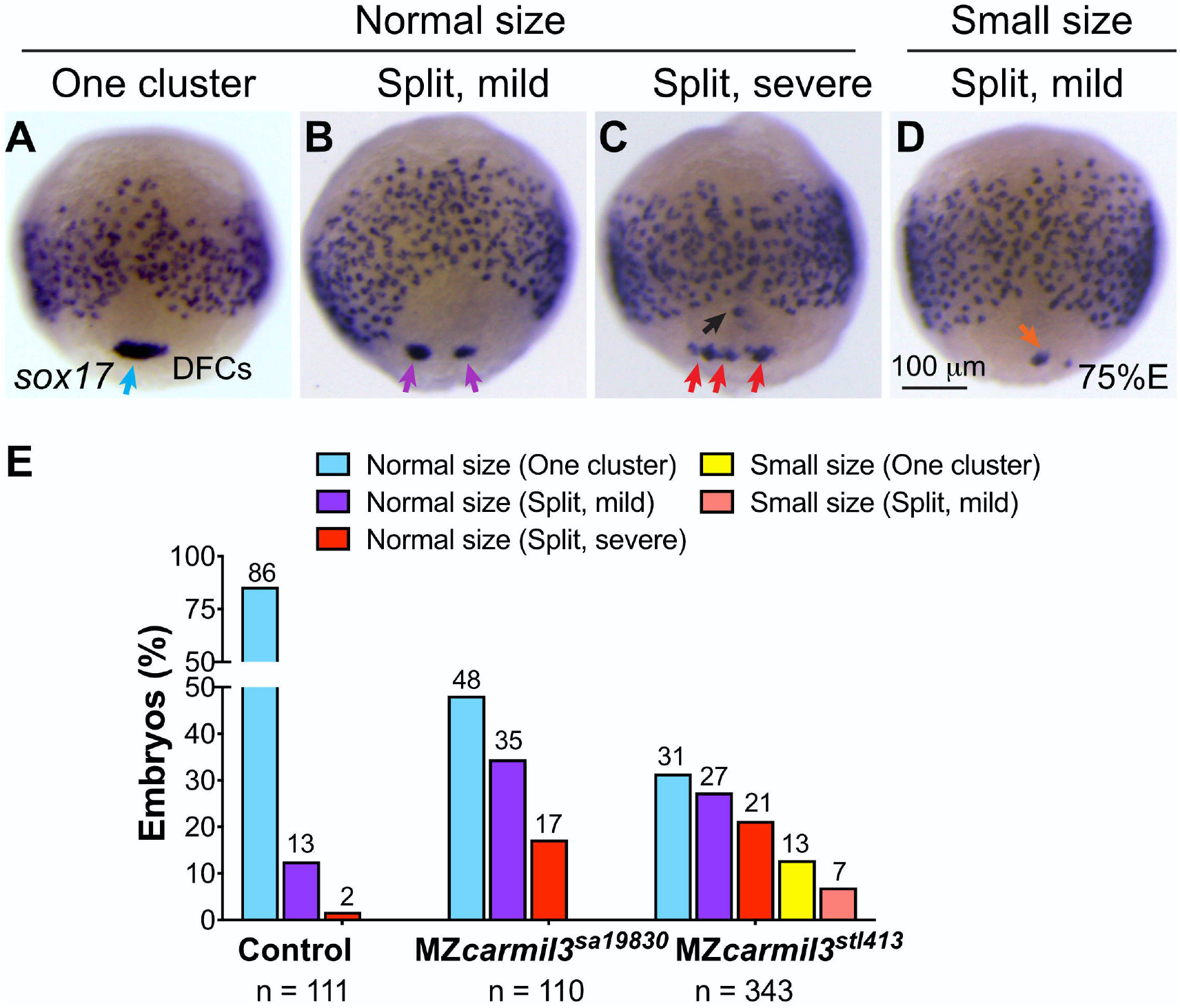
Patterns of dorsal forerunner cell (DFC) distribution in MZ*carmil3*^*sa19830*^ mutant embryos compared to heterozygous +/ MZ*carmil3*^*sa19830*^ embryos. Panels A to D illustrate the patterns observed, with results quantified in panel E. In most embryos, DFCs form a single tight cluster of normal size (light blue arrow in panel A and light blue bars in panel E). In some embryos, DFCs display three patterns of defects: B) normal size with mild splitting (i.e. two clusters); C) normal size with severe splitting (more than two clusters), and D) small size with mild splitting. (E) Percentages of embryos displaying the patterns of DFC defects in both MZ*carmil3*^*sa19830*^ and MZ*carmil3*^*stl413*^ mutants. The percentage of embryos in each group is indicated above each bar, and the total number of embryos in each group (N) is listed underneath the labels on the abscissa.

### Morphology of the Kupffer’s Vesicle and Cilia

The cluster of DFCs that forms during gastrulation, undergoes a morphogenetic transition into an epithelial sac known as Kupffer’s Vesicle (KV), the ciliated organ that determines L/R asymmetry during zebrafish development (Gokey et al., 2016). Because we observed a defect in DFC aggregate formation, we examined the morphology of the KV and its cilia during segmentation (Fig. 4).

Imaging the KV in MZ*carmil3*^*stl413*^ mutant (*Tg[sox17:EGFP]*) embryos at the 6-somite stage (Fig. 4A,B,B’), revealed that their KV during early segmentation was significantly smaller than in control WT embryos (Fig. 4C). To visualize the epithelial KV morphology during later development, we stained embryos fixed at the stage of 10-14 somites with antibodies against the epithelial marker atypical PKC (aPKC) (Amack et al., 2007) and acetylated tubulin (Ac-tubulin) (Piperno and Fuller, 1985), resolving the cilia as individual structures, allowing us to count their number and measure their length. In these images, the hollow empty center of the KV appears as a relatively dark unstained region in the aPKC channel and as staining positive for cilia in the Ac-tubulin channel. In instances where the KV did not “inflate” (MZ*carmil3*^*sa19830*^ in Fig. 4D, bottom panel), an unstained region was not observed. As in earlier stages, the KV was often smaller or fragmented, resulting in a decreased KV area for both mutants (Fig. 4C and 4E). The decrease in KV area was statistically significant (Fig. 4E), and the magnitude of the decrease was sufficiently large that the values were at or below the value for the area of the KV at this stage found to be necessary for robust L/R patterning by Gokey and colleagues (Gokey et al., 2016), which is drawn as a dotted line at 1300 μm^2^ in Figure 4E.

We analyzed the cilia in greater detail, and found that the number of cilia per KV was decreased in both mutants (Fig. 4F). The decreases were substantial and significantly different from WT embryos. In Figure 4F, the dotted line at 30 cilia per KV corresponds to the value that was found to be necessary for robust L/R patterning by Sampaio and colleagues (Sampaio et al., 2014). In both mutants, the number of cilia per KV was below this threshold (Fig. 4F). In contrast, the length of the cilia did not differ in the mutant embryos compared to WT embryos (Fig. 4G), suggesting that Carmil3 is not required for cilia biogenesis per se, but instead plays a role in the size and/or morphology of the KV.

**Figure 4.**
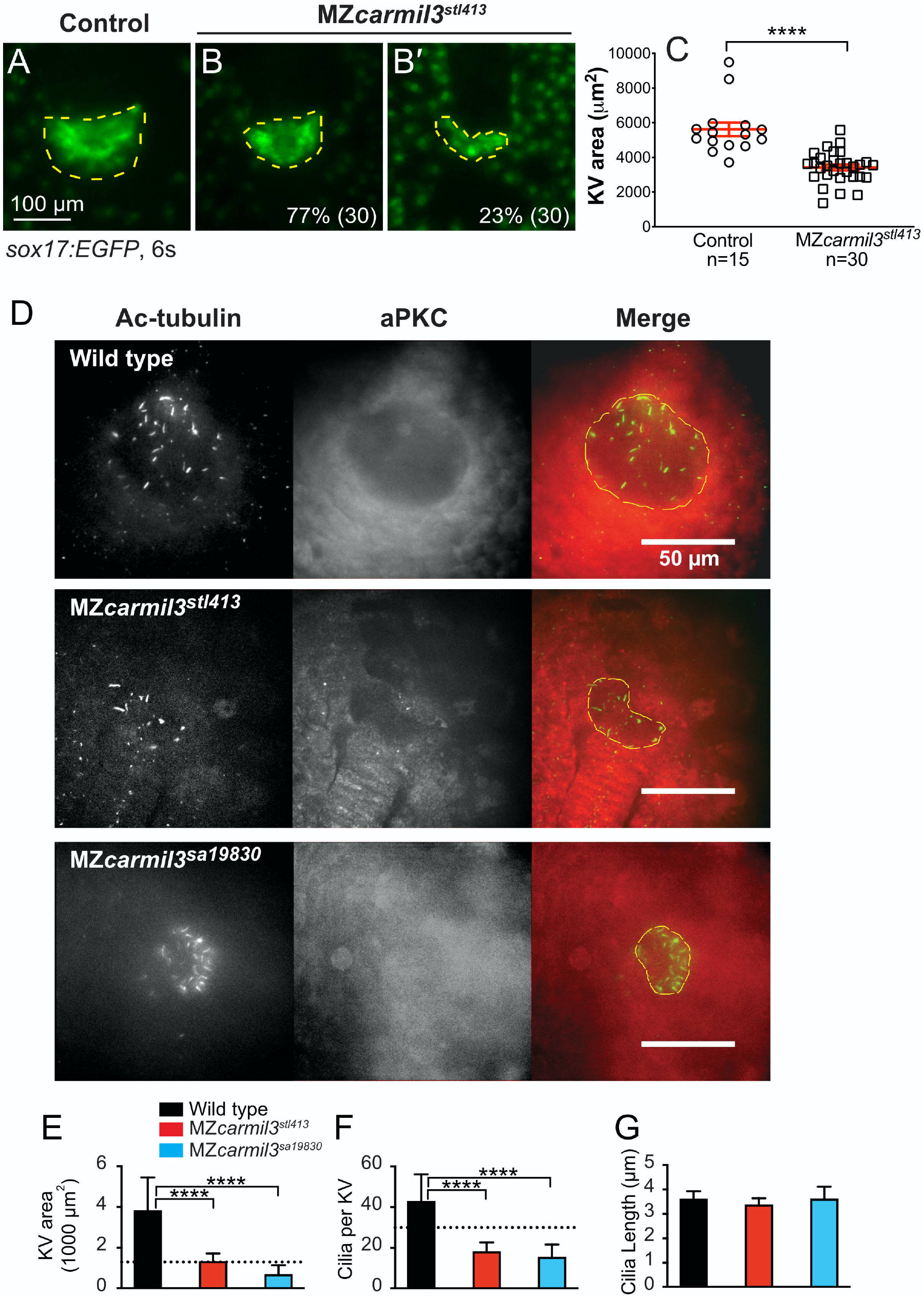
Morphology of Kupffer’s vesicle (KV) and cilia in *carmil3* mutant embryos. Panels A through C are from one analysis, based on wide-field fluorescence images showing the morphology of sox17-EGFP labelling of the KV in control (A) and MZ*carmil3*^*stl413*^ mutant (B-B’) embryos at the 6-somite stage. (C) Area of the KV in control and MZ*carmil3*^*stl413*^ mutant embryos. All the data points are shown. The red lines indicate the mean and one standard error of the mean. The results are statistically significant with a p value of < 0.0001. The number of embryos analyzed was 15 for control and 30 for mutant. In this set of experiments, the control was a heterozygous strain. Panels D through G are from a second experimental series with a different set of animals, based on confocal images of embryos at the 10-14 somite stage stained to visualize cilia and the KV, with antibodies to acetylated tubulin (Ac-tubulin, Green in Merge), and to atypical PKC (aPKC, Red in Merge) respectively. With anti-aPKC staining, the KV appears as a relatively dark area, owing to the absence of cells inside the vesicle. WT and two different *carmil3* mutant embryo lines are shown. The Merge panel shows examples of how the KV was outlined for calculation of area; the outline is based on both Ac-tubulin and aPKC images. Panels E through G are graphs of parameters quantified from the images, with the color scheme for WT and mutants as indicated. In each panel, the plotted values are the mean, and the error bars correspond to one standard deviation. E) Area of the KV. Horizontal dotted line corresponds to the value for area of KV found to be necessary for robust L/R patterning by Gokey and colleagues (Gokey et al., 2016). Values for the two mutants differ from the value for WT based on Student’s t-test (p<0.005 (actual=0.0006 and 0.0005)). F) Number of cilia per KV. Horizontal dotted line corresponds to the value for number of functional cilia per KV (30) found to be necessary for robust left / right patterning by Sampaio and colleagues (Sampaio et al., 2014). Values for the two mutants differ from the value for wild type based on Student’s t-test (p<0.005 (Actual=0.0001 and 0.0004)). G) Length of cilia. Values for the two mutant lines do not differ from the value for WT embryos based on Student’s t-test (p=0.3277 and >0.9999). Values of N as in panel E. Values of N (embryos counted) as follows: WT, 9; MZ*carmil3*^*stl413*^, 8; MZ*carmil3*^*sa19830*^, 4 for panels to G.

### Left-Right Patterning

Because we observed defects in formation of the KV, including its morphology, size and the number of cilia; we asked whether further development revealed defects in L/R asymmetry pathways and outcomes. Since the *carmil3* mutants displayed the KV area and number of cilia below the thresholds shown to be required for robust L/R patterning by previous studies (Gokey et al., 2016; Sampaio et al., 2014), we hypothesized that there would be defects in L/R patterning in these mutants.

First, we examined the pattern of expression of the lateral plate mesoderm marker *southpaw (spaw)* in embryos at the 18-20 somite stage (Fig. 5) (Long et al., 2003). Whereas WT embryos uniformly displayed *spaw* staining on the left, the two *carmil3* mutant lines showed substantial percentages of embryos with bilateral or right-sided *spaw* staining. Representative examples are shown in panel A of Figure 5, with quantitation in panel B and Table I. Results for the two mutant lines were similar to each other and differed significantly from results for WT embryos. These embryos were generated from the same set of mutant and WT animals as those used in the experiments illustrated in panels D-G of Figure 4, where the KV area and number of cilia per KV were below the critical threshold in the mutant. When a different generation of mutant animals was used in a separate set of experiments to generate MZ*carmil3*^*stl413*^ embryos, in which the KV area was not below the critical value illustrated in panels A-C of Figure 4, little or no defects in *spaw* staining distribution were observed (data not shown), consistent with the findings of Gokey and colleagues (Gokey et al., 2016), and Sampaio and colleagues (Sampaio et al., 2014) noted above.

**Table I.** Pattern of expression of *spaw* at 18-20 somites, comparing *carmil3* maternal zygotic (MZ) mutant embryos with WT embryos. The percentage of the total number of embryos with different patterns are listed, along with 95% confidence intervals (CI) and the number of embryos scored (N). Statistics, including confidence intervals, were calculated with GraphPad Prism.

|  | Phenotypic Pattern, Percent of Total (95% CI) |  |  |  |  |
| --- | --- | --- | --- | --- | --- |
| Genotype | Left | Right | Bilateral | Absent | N |
| Wild type | 98 (94–99) | 1 (0–3) | 2 (0–5) | 0 | 172 |
| MZ*carmil3*<sup>stl413</sup> | 54 (46–62) | 9 (5–15) | 26 (20–34) | 11 (7–17) | 137 |
| MZ*carmil3*<sup>sa19830</sup> | 53 (43–64) | 7 (3–14) | 31 (22–41) | 9 (5–17) | 88 |

**Figure 5.**
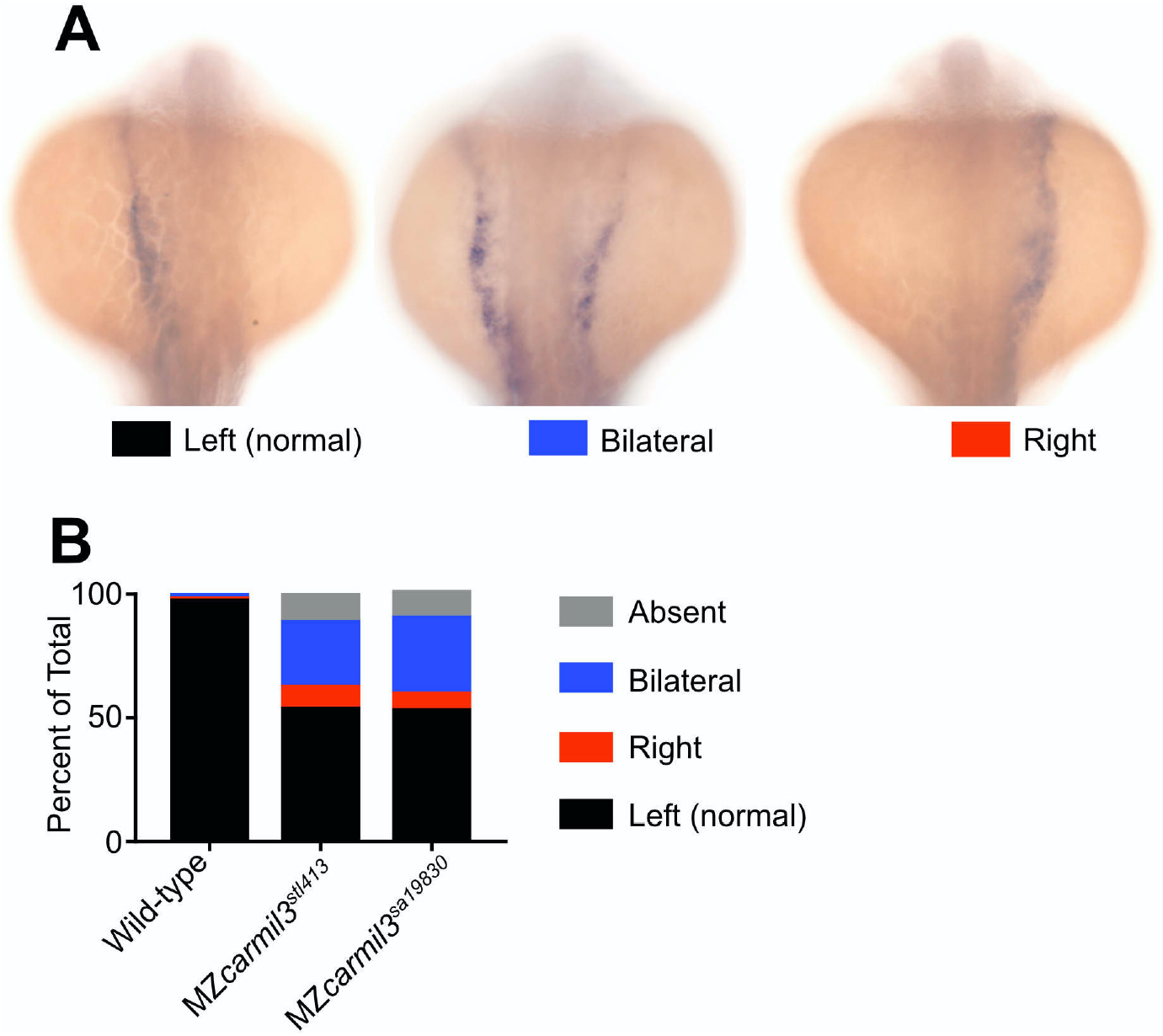
Patterns of *spaw* staining distribution at the 18-20 somite stage in *carmil3* mutant embryos compared with WT embryos. A. Representative images illustrate observed patterns of *spaw* staining, which is purple. B. Percentage of *spaw* staining patterns, comparing WT embryos with embryos of two different *carmil3* mutant lines, MZ*carmil3*^*stl413*^ and MZ*carmil3*^*sa19830*^. The color scheme is indicated in panel A, below the images. In cases scored as “absent,” no staining was observed. Values are listed in Table I.

*spaw* regulates FGF signaling (Neugebauer and Yost, 2014), and *spaw* mutants are defective in L/R asymmetry, as revealed by the position of the heart (Ahmad et al., 2004; Long et al., 2003). Therefore, we asked whether mutant embryos had a phenotype related to heart position. Indeed, mutant embryos at ~30 hpf displayed a defect in the L/R positioning of the heart (Fig. 6). In a noticeable and significant number of embryos, the heart was in the middle or on the right side of the embryo, based on staining for cardiac myosin light chain (*myl7*), for which representative examples are shown in panel A of Figure 6. A combined count of embryos with altered heart position showed defects of 6 - 9% for each of the two mutants and a value less than 1% observed in WT embryos (Fig. 6, Panel B). The defects in the two mutant lines did not differ from each other by a statistically significant degree, and the differences between the two mutant lines and the WT line was highly significant in both cases.

**Figure 6.**
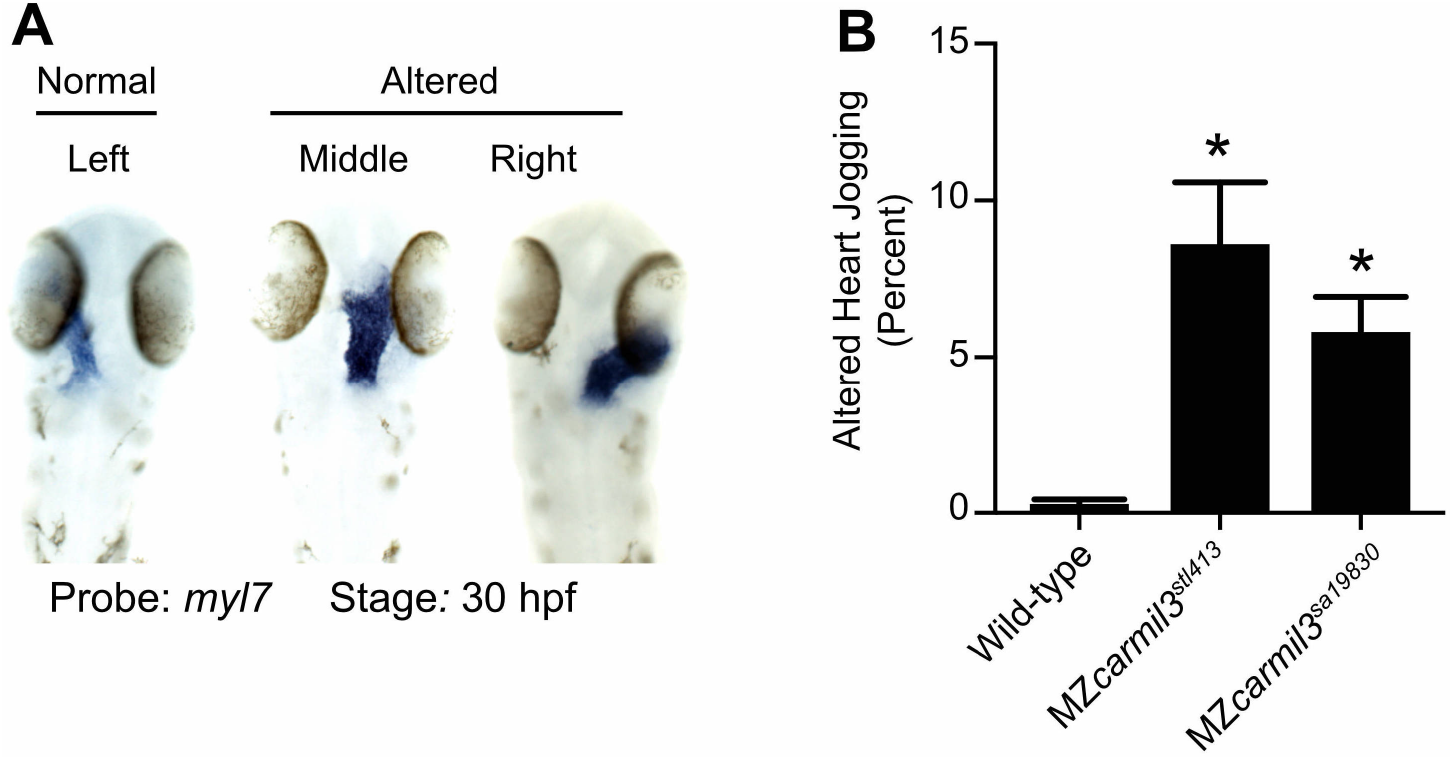
Heart position in MZ*carmil3* mutant embryos compared to WT embryos at ~30 hpf, assessed by staining for transcripts of *myl7*, the gene encoding cardiac myosin light chain, in blue. A. Representative images illustrate observed patterns. B. Quantification of the sum of the two altered patterns, termed “heart jogging,” in two different MZ*carmil3* mutant embryo lines compared with WT embryos. WT embryos displayed the abnormal (to the right or center) heart jogging phenotype 0.25% of the time (N=784). This value was significantly higher in the mutant embryos: 8.6% for MZ*carmil3*^*stl413*^ (N=198) and 5.8% for MZ*carmil3*^*sa19830*^ (N=415). Error bars indicates standard error of proportion. P values were calculated from Fisher’s exact test calculated with GraphPad Prism. Asterisk indicate that for both mutants, compared with WT, p values were <0.0001.

As above for *spaw* staining, these data for heart position are from a set of animals and experiments where the KV area and number of cilia per KV in the mutant were below the critical threshold (Fig. 4 D-G). In a separate set of experiments, using MZ*carmil3*^*stl413*^ mutants, in which the KV area was not below the critical value (Fig. 4 A-C), little or no defects in heart position were observed (data not shown). Therefore, the penetrance and expressivity of the KV phenotype in MZ*carmil3*^*stl413*^ mutants correlated well with those of the *spaw* asymmetric staining defect during mid-segmentation stages and the L/R heart positioning defect at the end of embryogenesis, consistent with the findings of Gokey and colleagues (Gokey et al., 2016) and Sampaio and colleagues (Sampaio et al., 2014).

## Discussion

We investigated the role of the actin assembly regulator, CARMIL3, in cell migration and morphogenesis during early zebrafish development. First, we confirmed that in actin polymerization assays, human CARMIL3 and zebrafish Carmil3 interact with and regulate the activity of capping protein with respect to actin polymerization. More important, we discovered that Carmil3 is required for normal migration of endodermal cells and the aggregation of dorsal forerunner cells (DFCs) during zebrafish gastrulation. Impaired aggregation of DFCs led to defects in the formation of the KV in terms of its shape and size. These KV defects were associated with decreased numbers of cilia present in mutant KVs; as a consequence, L/R asymmetry was impaired, manifested by malposition of the marker *spaw* during segmentation and later in the position of the zebrafish heart tube. These KV and L/R phenotypes displayed variable but correlated penetrance and expressivity.

The two *carmil3* mutant alleles we report here were generated by reverse genetic approaches: *carmil3*^*sa19830*^ by TILLING for ENU-induced nonsense mutations (Kettleborough et al., 2013) and *carmil*^*stl413*^ by deploying TAL endonucleases (Boch et al., 2009; Moscou and Bogdanove, 2009). Both are premature stop codons predicted to produce truncated proteins and thus should represent strong or null alleles. Consistent with that view, the phenotypes are recessive. However, our study does not include an analysis of the protein products that would test this view definitively. The observation that only maternal zygotic but not zygotic *carmil3* mutants presented gastrulation phenotypes, is consistent with known significant maternal *carmil3* expression, which is likely to compensate for the lack of zygotic expression during gastrulation (Solnica-Krezel, 2020; Stark and Cooper, 2015). The lack of full penetrance for the later phenotypes indicates a substantial level of robustness in the processes of cell migration and morphogenesis, consistent with observations for many other genes regulating early embryogenesis (Chen et al., 2018; Kelly et al., 2000; Li-Villarreal et al., 2015; Solnica-Krezel and Driever, 2001). CARMILs are encoded by three conserved genes in vertebrates, including zebrafish, so redundant and overlapping functions contributed by CARMIL1 or CARMIL2 may also account for the variable level of penetrance seen for downstream phenotypes. In addition, as the two *carmil3* mutant alleles are nonsense mutations, the observed phenotypes could represent only partial loss-of-function and variability due to genetic compensation triggered by RNA degradation (El-Brolosy et al., 2019; Ma et al., 2019).

Our results provide new information about the function of CARMIL3, especially in the *in vivo* context of a whole vertebrate organism during the process of embryogenesis. Previous studies on CARMIL3 have used cell culture and mouse tumor models to uncover roles in neuronal synapse formation and cancer cell migration, based on actin assembly (Hsu et al., 2011; Lanier et al., 2016; Spence et al., 2019; Wang et al., 2020).

Previous studies, with human cultured cells that express both CARMIL1 and CARMIL2, found overlapping but distinct functions for the two proteins, based on subcellular localization and knockdown phenotypes (Liang et al., 2009; Stark et al., 2017). While the early phenotypes observed here display strong penetrance, the later phenotypes are far less penetrant, raising the question of compensatory and overlapping function among the CARMIL-encoding genes. In support of the view, preliminary observations in our laboratories have revealed stronger L/R asymmetry phenotypes in double mutant zebrafish embryos that carry mutations in genes for both CARMIL2 and CARMIL3 (Stark, Solnica-Krezel and Cooper, 2019, unpublished); these observations will merit further study in the future.

We discovered that Carmil3 is important for the migration of endodermal cells during zebrafish gastrulation. Mutant endodermal cells had both reduced overall motility as well as reduced persistence. Recent work demonstrated cultured cells lacking CARMIL3 had reduced migration in the classical scratch assay and in trans-well migration. KO cells were less polarized, had less polymerized actin and fewer focal adhesions (Wang et al., 2020). Interestingly, the Rac-specific guanine nucleotide exchange factor, Prex1, was implicated in regulation of Nodal-dependent actin dynamics and random endoderm cell motility during gastrulation (Woo et al., 2012). Future experiments will determine if endodermal migration defects in MZ*carmil3* mutants are associated with abnormal actin organization and/or focal adhesion formation.

We found that Carmil3 is important for the aggregation of DFCs during gastrulation. DFCs typically migrate vegetalward ahead of the germ layers in one cluster of cells. Occasionally a few cells may separate from the cluster. DFC coalescence and migration as a cohesive cluster is strongly dependent on cell-cell adhesion. *cdh1/*E-cadherin (*halfbaked/volcano)* mutants in which cell adhesion is reduced, exhibit delayed epiboly of all germ layers and frequently fragmented DFC clusters (Shimizu et al., 2005; Solnica-Krezel et al., 1996). A recent study showed loss of CARMIL3 in cultured cells inhibits cadherin based adhesion through transcriptional downregulation of epidermal type gene expression (Wang et al., 2020). Such a mechanism might account for the reduced adhesion of DFC *in vivo*. We observed small and malformed KV, likely the direct result of the smaller DFC clusters. However, we cannot exclude a more direct role of Carmil3 in KV morphogenesis.

CARMILs appear to regulate actin via direct biochemical interactions with and effects on the actin-capping properties of CP (Edwards et al., 2013; Lanier et al., 2015; Stark et al., 2017). Indeed, CP itself is known to have an important role in morphogenesis. Mutations in humans and in zebrafish of the gene encoding the beta subunit of CP, known as *capzb* in zebrafish, were found to cause craniofacial and muscle developmental defects, with effects on cell morphology, cell differentiation and neural crest migration (Mukherjee et al., 2016). In addition, the same study found that *capzb* overexpression produced embryonic lethality.

Among regulators of CP, CARMILs are only one family of proteins with CPI motifs. Among other CPI-motif protein families, which are unrelated to each other outside of their CPI motifs, several have been shown to have roles in actin-based processes of development. The CPI-motif protein CKIP-1 is important for myoblast fusion in mammalian and zebrafish systems (Baas et al., 2012). CapZIP, known as *duboraya/dub* in zebrafish, is important in zebrafish development for actin organization in cells lining the KV, cilia formation in the KV, and L/R asymmetry (Oishi et al., 2006). CD2AP, encoded by *cd2ap* in zebrafish, is important for the development and function of the kidney glomerulus, in mammals and zebrafish (Hentschel et al., 2007; Tossidou et al., 2019). Zebrafish CIN85, a homologue of CD2AP encoded by the gene *sh3kb1*, is also important for glomerular podocyte function (Teng et al., 2016), and it has a role in the formation and maintenance of the vascular lumen (Zhao and Lin, 2013). Finally, the twinfilin family of CPI-motif proteins, encode by four genes in zebrafish - *twf1a, twf1b, twf2a* and *twf2b*, has not been as well-studied, but may also affect actin assembly, as may the WASHCAP / Fam21 family of CPI-motif proteins, encoded by *washc2c* in zebrafish.

The *carmil3* mutant lines generated here provide a valuable tool for studying the regulation of actin dynamics during endoderm migration and KV morphogenesis in the context of a developing vertebrate embryo, and for testing potential functional interactions with CARMIL3 interacting proteins.

## Acknowledgments

We are grateful to members of our laboratories for advice and assistance. We are grateful for the activities and support of the Washington University Zebrafish Facility and the University of Iowa Zebrafish Facility. This research was supported by the following grants: NIH R35 GM118171 to JAC, and NIH R35 GM118179 to LSK.

